# Metabolomic Profiles and Pathways in Osteoarthritic Human Cartilage: A Comparative Analysis with Healthy Cartilage

**DOI:** 10.1101/2024.01.25.577269

**Authors:** Hope D. Welhaven, Avery H. Welfley, Priyanka Brahmachary, Annika R. Bergstrom, Eden Houske, Matthew Glimm, Brian Bothner, Alyssa K. Hahn, Ronald K. June

**Affiliations:** Department of Chemistry & Biochemistry, Montana State University, Bozeman MT; Department of Mechanical & Industrial Engineering, Montana State University, Bozeman MT; Department of Chemical & Biological Engineering, Villanova University, Villanova, PA; Department of Biological and Environmental Sciences, Carroll College, Helena, MT

**Keywords:** Osteoarthritis, Cartilage, Metabolomics, Mass Spectrometry

## Abstract

**Objective:** Osteoarthritis (OA) is a chronic joint disease with heterogenous metabolic pathology. To gain insight into OA-related metabolism, healthy and end-stage osteoarthritic cartilage were compared metabolically to uncover disease-associated profiles, classify OA-specific metabolic endotypes, and identify targets for intervention for the diverse populations of individuals affected by OA.

**Design:** Femoral head cartilage (n=35) from osteoarthritis patients were collected post-total joint arthroplasty. Healthy cartilage (n=11) was obtained from a tissue bank. Metabolites from all cartilage samples were extracted and analyzed using liquid chromatography-mass spectrometry metabolomic profiling. Additionally, cartilage extracts were pooled and underwent fragmentation analysis for biochemical identification of metabolites.

**Results:** Specific metabolites and metabolic pathways, including lipid- and amino acid pathways, were differentially regulated between osteoarthritis-derived and healthy cartilage. The detected alterations of amino acids and lipids highlight key differences in bioenergetic resources, matrix homeostasis, and mitochondrial alterations in osteoarthritis-derived cartilage compared to healthy. Moreover, metabolomic profiles of osteoarthritic cartilage separated into four distinct endotypes highlighting the heterogenous nature of OA metabolism and diverse landscape within the joint between patients.

**Conclusions:** The results of this study demonstrate that human cartilage has distinct metabolomic profiles between healthy and end-stage osteoarthritis patients. By taking a comprehensive approach to assess metabolic differences between healthy and osteoarthritic cartilage, and within osteoarthritic cartilage alone, several metabolic pathways with distinct regulation patterns were detected. Additional investigation may lead to the identification of metabolites that may serve as valuable indicators of disease status or potential therapeutic targets.

## 1. INTRODUCTION

Osteoarthritis (OA) is the leading cause of disability worldwide. Since 1999, the number of global cases has increased by an astonishing 113%, equating to ∼528 million individuals affected in 2019^1, 2^. In the United States alone, 32.5 million adults have OA, costing $185 billion annually^3–6^. At the heart of OA’s insidious progression lies the gradual breakdown of articular cartilage (AC) and other joint tissues. The imbalanced activity between matrix anabolism and catabolism contributes to the observed changes in AC, other tissues, and fluids affected by OA (*i.e.*, underlying bone, synovium, synovial fluid). Previous studies have examined altered metabolism in various OA-associated tissues and their cell types, such as chondrocytes, to investigate disease-associated metabolic activity^7–9^. However, significant limitations of many studies are that they are performed *in vitro* and/or lack healthy human controls, thereby hindering a complete understanding of the role metabolism plays in OA development.

Moreover, the complex nature of OA can manifest differently between individuals. Specifically, symptom severity, rate of progression, response to treatment, pain perception as well as other factors can vary from person to person^10–12^. Therefore, a “one-size-fits-all” approach to the treatment and prevention of OA is limited. More recent studies describe OA as a group of symptoms encompassing multiple distinct phenotypes and endotypes rather than a single disease^13^. Previously, phenotype was defined as a single or collection of disease characteristics that explain differences between patients and their outcomes, such as symptom severity^11–13^.

Conversely, endotype is defined functionally and pathologically by a molecular mechanism noting that different mechanisms can lead to the same manifestation, such as end-stage OA^14^. Examining OA phenotypes and endotypes may shed light on the epidemiological origins and development of OA, unveil biomarkers, and lead to targeted interventions for sub-populations of OA individuals, all of which have potential to improve patient outcomes.

Metabolomics, the study of small molecule intermediates called metabolites^15^, is advantageous for generating and investigating OA metabolic endotypes because it detects thousands of metabolites. This enables the generation of biochemical signatures that represent the overall physiological state of the tissue. To our knowledge, two prior studies used a similar approach to examine synovial fluid metabolism from OA individuals. Here, researchers characterized different regulation patterns, or endotypes, based on detected differences in biochemical signatures between healthy and OA individuals^16, 17^. However, this same approach has yet to be applied to osteoarthritic cartilage. To begin filling these gaps in knowledge, we compared the metabolome of radiography-confirmed end-stage OA cartilage (Kellgran-Lawrence grades III and IV) with healthy cartilage using liquid chromatography-mass spectrometry (LC-MS) metabolomics.

Thus, the primary objective of this study was to identify disease-associated OA metabolomic profiles to shed light on the pathological mechanisms underlying OA. The secondary objective was to examine and classify endotypes of OA. Furthermore, we used tandem LC-MS (LC-MS/MS) for biochemical identification of key metabolites. This has the potential to identify novel biomarkers and drug targets to slow, halt, or reverse OA progression. With this approach, we aimed to uncover specific metabolic endotypes and metabolite identities to serve as potential indicators of disease status or therapeutic intervention across sub-populations of OA individuals.

## 2. METHODS

### 2.1 Articular cartilage sample obtainment

Under IRB approval, 35 femoral heads from end-stage OA patients were obtained following total joint arthroplasty from local musculoskeletal clinics. Partial patient information including age, sex, age, and BMI was provided (Supplemental Table 1). However, radiographic scans were not obtained due to IRB approval only permitting partial patient information to be shared. Post-mortem cartilage samples obtained from donors without joint disease (Articular Engineering) to serve as healthy controls for comparison.

### 2.2 Metabolite extraction and mass spectrometry analysis

Cartilage samples were shaved from the femoral head prior to metabolite extraction. All cartilage samples (n=3<u>5</u> OA, n=11 healthy) were extracted using a previously established protocol. All cartilage samples were weighed prior to extraction for normalization purposes. Notably, the weights of healthy cartilage consistently measured (100 mg), while OA cartilage exhibited variation due to their origin from end-stage OA patients, with differences in the extent of cartilage integrity among individuals. Next, cartilage shavings were submerged in 3:1 methanol:water and homogenized using a tissue homogenizer (SPEX Sample Prep 1200 GenoLyte, Fisher Scientific). Homogenization included 15 cycles of homogenizing for 20 seconds and resting for 2 minutes. Next, samples were briefly vortexed and stored at −20°C overnight to promote protein precipitation. The following day, samples were vortexed again, centrifuged for 10 minutes at 16100 *x g* at 4°C, and supernatants were collected and dried via vacuum concentration. Dried supernatants were then resuspended with 1:1 acetonitrile:water, stored at −20°C for 30 minutes, then centrifuged again for 10 minutes at 16100 *x g* at 4°C. Similarly, supernatants were dried via vacuum concentration and then prepped for liquid chromatography-mass spectrometry (LC-MS) by resuspending with 1:1 acetonitrile:water. Additionally, pooled samples (n=4) were generated by randomly pooling 5 µL from each of 10 individual extracts.

All samples, including extracted cartilage and pools, underwent LC-MS analysis as previously described^18^. In brief, an Aquity UPLC Plus interfaced through an electrospray ionization source to a Waters Synapt XS was used. A Cogent Diamond Hydride HILIC column (150 x 2.1 mm) at a flow rate of 0.400 µL/min was used to separate metabolites over a 19-minute elution gradient (A = 95/5% water/acetonitrile, B = 95/5% acetonitrile:water). Every 10 injections, quality control blanks containing mass spectrometry-grade water were injected to minimize spectral drift and assess LC-MS performance. Cartilage extracts underwent standard LC-MS, whereas pooled samples underwent liquid chromatography tandem mass spectrometry (LC-MS/MS) at a constant high energy ramp of 30-50V for secondary ionization to derive metabolite identifications.

### 2.3 Statistical and metabolomic profiling

LC-MS data, consisting of mass-to-charge ratios (m/z), relative metabolite abundance, and retention time, were processed using MSConvert^19^ and XCMS^20^. Prior to data analysis, metabolites associated with each cartilage sample were normalized by the pre-extraction recorded cartilage shaving weight. Previously established analysis pipelines were used^18, 21^, and executed in MetaboAnalyst^22^ where data underwent interquartile range normalization, log-transformation, and autoscaling (mean-centered/standard deviation of each metabolite feature). In brief, hierarchical clustering analysis (HCA), principal component analysis (PCA), and partial least squares-discriminant analysis (PLS-DA) were used to visualize dissimilarities in metabolomic profiles between healthy and OA cartilage, as well as examine OA endotypes. T-test, fold change, and volcano plots were used to assess significance and magnitude of change. Moreover, these populations of metabolite features were differentially regulated between groups and identified by these tests underwent pathway enrichment analysis using MetaboAnalyst’s Functional Analysis feature, which utilizes the *mummichog* algorithm to predict networks of functional activity from metabolite features of interest. The pathway library, Human MFN, was used as the primary reference library to match metabolite features to putatively identified metabolites (mass tolerance: 5 parts per million (ppm), positive mode, version 1). Significance for pathway analyses, and all other statistical tests, were determined using a false discovery rate (FDR)-corrected significance level of p < 0.05.

### 2.4 Metabolite identification

A major hurdle in LC-MS-based metabolomics is metabolite identification^23^. To address this challenge, pooled samples were subjected to LC-MS/MS involving fragmentation allowing analysis of parent and daughter fragment ions. These data were manually analyzed to confirm metabolite identifications as follows. Firstly, all LC-MS/MS data from pooled samples were imported, peak picked, and aligned using Progenesis QI (Nonlinear Dynamics, Newcastle UK, version 3.0). Utilization of Progenesis improves efficiency of identifications and uses a computational framework that allows for the exploration of thousands of putative metabolite identifications across various databases. Here, the Human Metabolome Database (HMDB)^24^ was utilized to compare theoretical fragmentation patterns to the acquired fragmentation patterns of parent and daughter ions. For a metabolite identity to be deemed valid and subsequently investigated manually, we required identities to receive a fragmentation score and overall progenesis score greater than 12 and 60 out of 100, respectively. These score criteria are based on mass error, isotope distribution, and retention time. Once identified metabolites were narrowed based on these set scores and parameters, they were matched against populations of LC-MS-based metabolite features distinguished by statistical analyses comparing OA and healthy cartilage, as well as OA endotypes. To minimize false identifications, a threshold of 10 ppm error between observed and Progenesis-identified metabolites was enforced.

## 3. RESULTS

### 3.1 Global metabolomic profiles of osteoarthritis and healthy cartilage unveil altered cellular mechanisms associated with disease

In total, 10,853 metabolite features were detected by LC-MS across all cartilage samples. To visualize and assess metabolomic differences between healthy and OA cartilage, we used unsupervised (HCA, PCA) and supervised (PLS-DA) multivariate tests. HCA, visualized by a dendrogram and measured using Euclidean distance, displayed clear separation of healthy and OA cartilage (Figure 1A). A similar trend was observed when using PCA and PLS-DA where near perfect separation of groups was displayed demonstrating metabolomic profiles reflective of the disease status of cartilage (Figure 1B-C).

**Figure 1.**
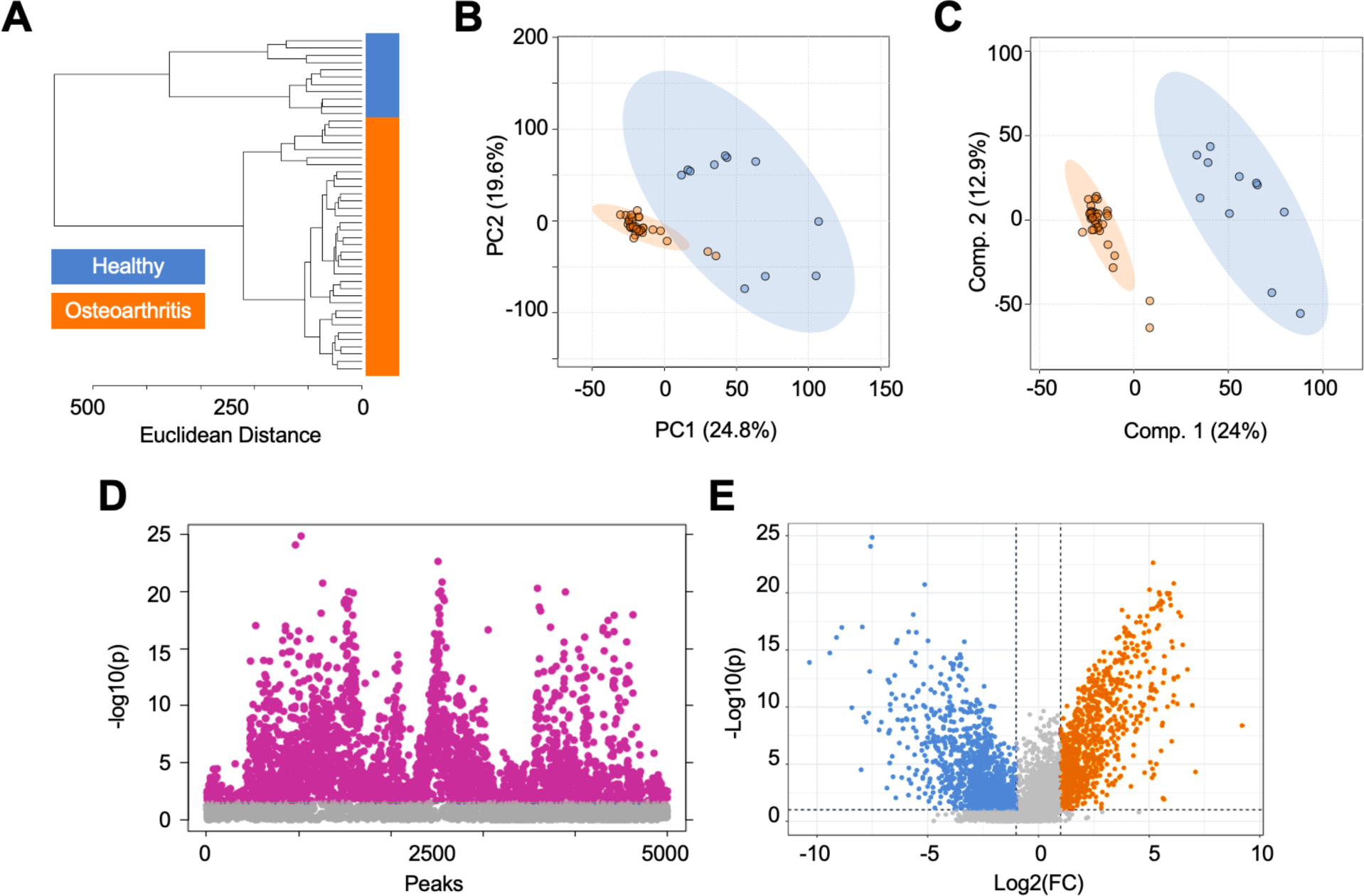
Metabolomic profiles of human cartilage from healthy and osteoarthritis patients are metabolically distinct. (A) Hierarchical clustering analysis finds that healthy and osteoarthritic cartilage samples cluster separate from each other. (B) Principal component analysis, an unsupervised test, finds minimal overlap with principal components 1 and 2 accounting for 44.4% of the variability in the dataset. (C) Partial least squares-discriminant analysis, a supervised test, shows complete separation of healthy and diseased cartilage samples with components 1 and 2 accounting for 36.9% of the variability in the dataset. (D) Volcano plot analysis, using fold change and statistical significance, distinguished differentially regulated metabolites between healthy and diseased cartilage. Specifically, 1,010 metabolite features were higher in abundance in diseased cartilage compared to healthy cartilage (log2(FC) > 2, p < 0.05), whereas 1,399 were higher in abundance in healthy cartilage compared to diseased cartilage (log2(FC) < −2, p < 0.05). (E) T-test analysis detected 2,842 metabolite features with a false discovery rate adjusted p-value less than 0.05. Orange = osteoarthritis. Blue = healthy.

Next, t-test and volcano plot analyses were performed to distinguish dysregulated populations of metabolite features between healthy and OA cartilage. Populations distinguished by both analyses were then analyzed using MetaboAnalyst’s Functional Analysis feature to find biological pathways that differ in regulation across groups. Volcano plot analysis found 987 metabolite features that were higher in abundance in OA cartilage compared to healthy (Figure 1D). These metabolite features mapped to numerous lipid-related pathways (omega 3 & 6 fatty acid metabolism, fatty acid activation & oxidation, polyunsaturated and saturated fatty acid beta-oxidation, glycosphingolipid metabolism), the carnitine shuttle, leukotriene metabolism, and others (Table 1, Supplemental Table 2). Conversely, volcano plot also found 1,193 metabolite features that were higher in abundance in healthy cartilage compared to OA cartilage (Figure 1D). These features mapped to the urea cycle, purine metabolism, glycerophospholipid metabolism, vitamin metabolism (K, E), squalene and cholesterol biosynthesis, aminosugar metabolism, and various amino acid metabolic pathways (methionine, cysteine, histidine, glycine, serine, alanine, threonine, tryptophan) (Table 1, Supplemental Table 2). T-test distinguished 2,842 metabolite features that were significantly dysregulated between groups (FDR p < 0.05) (Figure 1E).

**Table 1.**
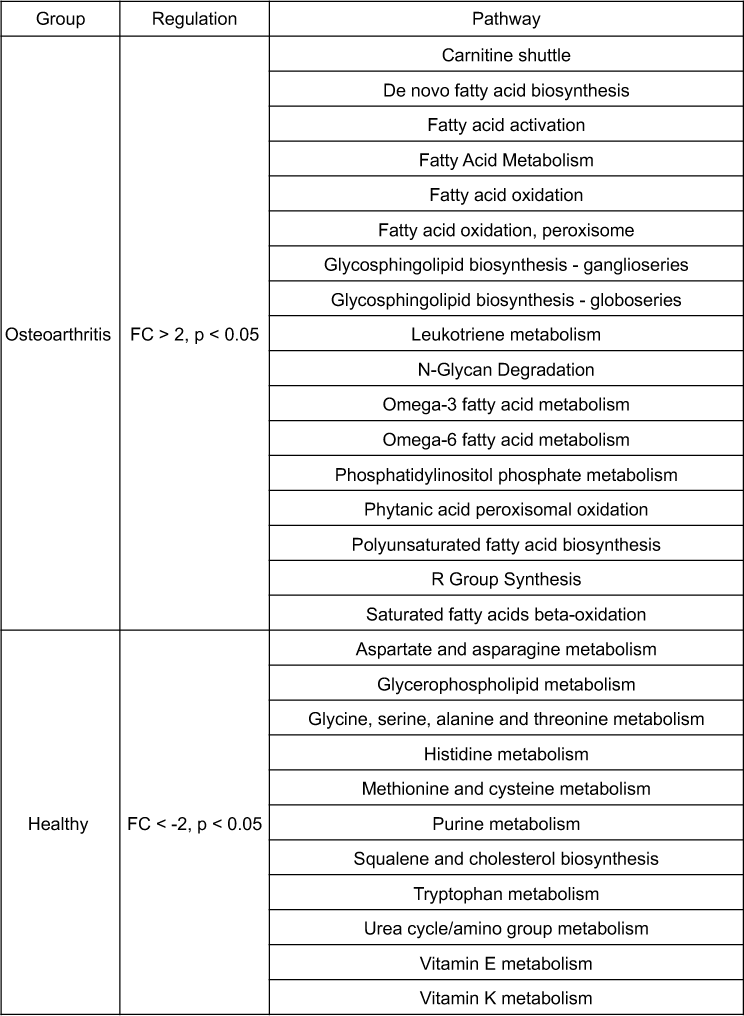
Metabolic pathways associated with healthy and diseased cartilage identified by volcano plot analyses. All reported pathways have a FDR-corrected significance level < 0.05.

Additionally, features distinguished by volcano plot (Supplemental Table 3) and t-test (Supplemental Table 4) were matched to putative identifications made using LC-MS/MS to unveil metabolic indicators of disease. Putatively identified metabolites that were statistically significant in both t-test and volcano plot analyses and were higher in healthy cartilage compared to OA cartilage included N-acetyl-leukotriene E4, demethylphylloquinone, 7C-aglycone, androsterone sulfate, and others (Supplemental Figure 1A). The majority of identified metabolites distinguished by these analyses were higher in abundance in healthy cartilage, with the exception of guanidinoethyl methyl phosphate, cervonyl carnitine, erythromycin propionate, and glycocholic acid (Supplemental Figure 1B). Collectively, these findings unveil specific metabolites and metabolic pathways that show altered cellular mechanisms of OA and reflect disease status of cartilage.

### 3.2 Endotype characterization supports heterogenous nature of osteoarthritis

To examine the heterogenous nature of OA metabolism and better understand differences in the diverse metabolic landscape within OA, we examined metabolomic endotypes. HCA (Figure 2A) and ensemble clustering (Supplemental Figure 2) were utilized to identify OA endotypes across all cartilage samples. The application of both methods aimed to minimize subjectivity in delineating OA cartilage endotypes. These analyses unveiled four distinct endotypes of OA participants. Considering patient-specific factors like age, the ratio of males to females, and overall number of participants within each endotypes, we found no clear pattern related to participant demographics that correlated with these four endotypes (Supplemental Table 1).

**Figure 2.**
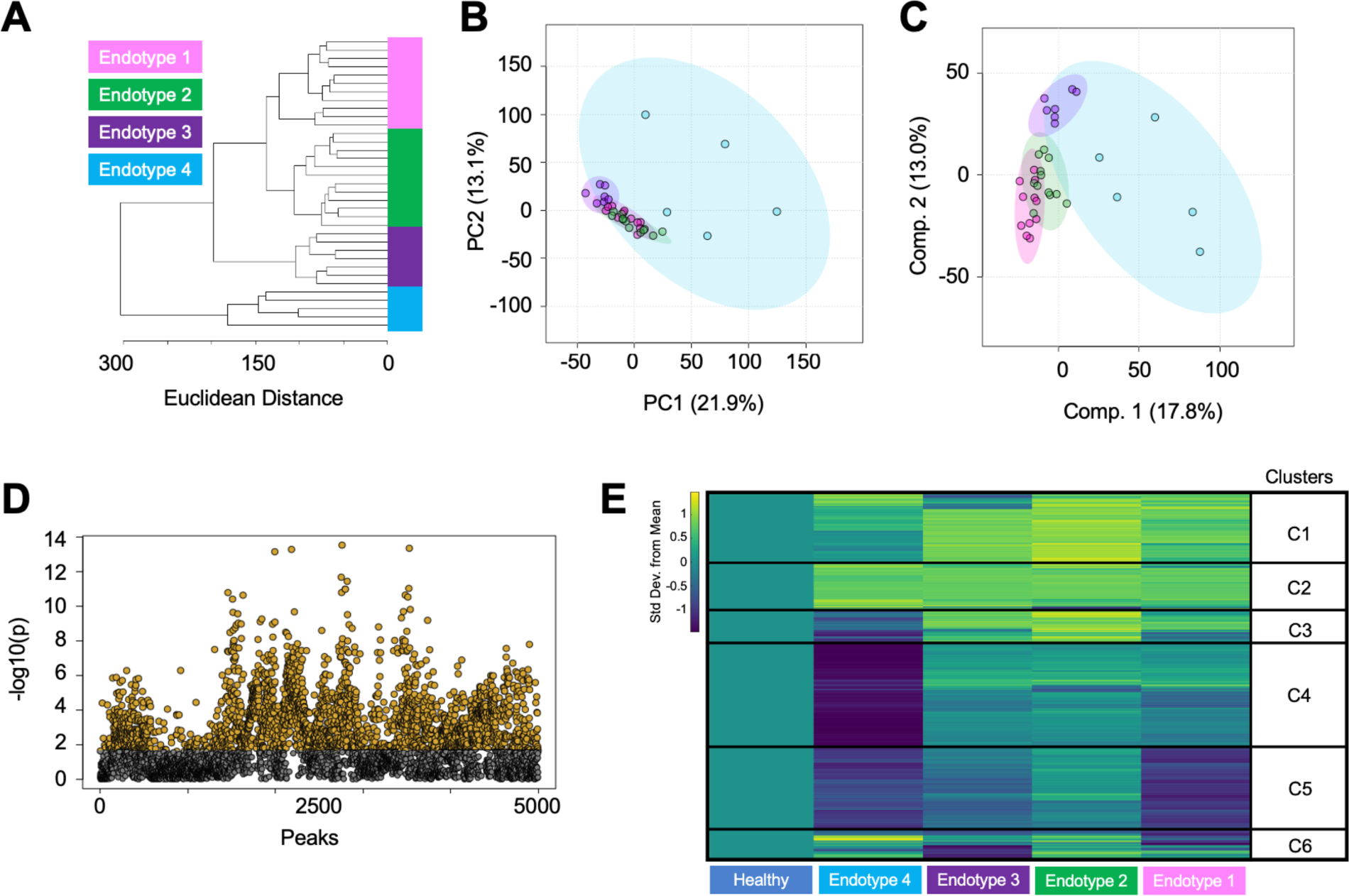
Metabolomic assessment of osteoarthritic cartilage classifies unique patient endotypes. (A) Hierarchical clustering of cartilage from patients with osteoarthritis cluster into 4 distinct endotypes. (B) Endotypes were further examined and visualized by principal component analysis. Principal components 1 and 2 account for 35% of the variability in the dataset and showed moderate overlap of osteoarthritis endotypes. (C) Partial least squares-discriminant analysis slightly refines separation of groups with components 1 and 2 accounting for 30.8% of the variability in the dataset. (D) ANOVA analysis detected 2,508 metabolite features with a false discovery rate adjusted p-value less than 0.05. (E) Metabolite features distinguished by ANOVA were visualized using a median metabolite intensity heatmap analysis where osteoarthritis endotypes were normalized to healthy cartilage. Clusters of co-regulated metabolite features (C1-C6) were then subjected to pathway analyses to pinpoint distinct metabolomic endotypes across osteoarthritic cartilage. Columns represent endotype and rows indicate metabolite features. Warmer colors (yellow) indicate higher metabolite abundance, whereas cooler colors (blue) indicate lower metabolite abundance. Endotype colors correspond to: pink – Endotype 1; green – Endotype 2; purple – Endotype 3; blue – Endotype 4.

Once endotypes were distinguished and patient-specific factors were examined, we used PCA and PLS-DA to gain additional insight into these endotypes. This revealed limited overlap between endotype groups (Figure 2B-C). Notably, endotype 4 exhibited considerable variability portrayed by a substantial ellipse. In contrast, endotypes 1-3 showed a closer metabolomic resemblance where the proximity of smaller and tighter clustered ellipses were observed.

Subsequent ANOVA analysis identified 2,506 metabolite features that were significantly different between endotypes with FDR-corrected p < 0.05 (Figure 2D). This subset of features was then matched to putative identifications made using LC-MS/MS data. Identifications consisted of lipid and lipid-like metabolites including cervonyl carnitine, lucidenic acid A, 6-Epi-7-isocucurbic acid glucoside, various phosphatidylcholine species, and others (Supplemental Figure 3A, Supplemental Table 6). Additionally, metabolites related to arachidonic acid and leukotriene metabolism were identified including arachidonic acid, panaxydol linoleate, leukotriene F4, and N-acetyl-leukotriene E4 (Supplemental Figure 3B, Supplemental Table 5).

Additionally, this subset of statistically significant metabolite features were further examined using a median metabolite intensity heatmap analysis normalized to healthy cartilage to find dissimilarities in metabolic regulation across the four OA endotype groups compared to healthy cartilage (Figure 2E). Metabolite features within clusters 1 and 2, which exhibited higher abundances across endotypes compared to healthy cartilage, mapped to 16 statistically significant pathways including leukotriene metabolism, selenoamino acid metabolism, the carnitine shuttle, and numerous lipid-related pathways (Table 2, Supplemental Table 6). Clusters 3 and 4, comprised of metabolites that were lowest in endotype 4 but highest in other endotypes, mapped to 16 statistically significant pathways including vitamin A metabolism, phytanic acid peroxisomal oxidation, lysine and tyrosine metabolism, keratan sulfate degradation, N-glycan degradation, lineolate metabolism, butanoate metabolism, and various lipid-related pathways. Metabolites composing cluster 5 were relatively lower across endotypes compared to healthy samples which mapped to 7 statistically significant pathways including purine metabolism, leukotriene metabolism, urea cycle, aminosugar metabolism, and various amino acid pathways (methionine, cysteine, tryptophan, aspartate, asparagine). Lastly, cluster 6 consisted of metabolites with mixed regulation patterns across endotypes; however, no statistically significant pathways were detected. All pathways reported had an FDR-corrected p < 0.05. Collectively, these findings underscore the heterogenous nature of OA metabolism among those with OA and provide compelling evidence to support the diverse landscape of metabolic regulation associated with this disease.

**Table 2.** Metabolic pathways associated with osteoarthritis endotypes classified by median intensity heatmap analysis. All pathways reported have a FDR-corrected significance level < 0.05. Clusters defined in. **Figure 2E**.

| Cluster | Pathway |
| --- | --- |
| 1 | Fatty acid activation |
| 1 | Saturated fatty acids beta-oxidation |
| 1 | De novo fatty acid biosynthesis |
| 1 | Fatty Acid Metabolism |
| 1 | Omega-6 fatty acid metabolism |
| 1 | Carnitine shuttle |
| 1 | R Group Synthesis |
| 1 | Fatty acid oxidation |
| 1 | Fatty acid oxidation, peroxisome |
| 1 | Leukotriene metabolism |
| 2 | Fatty acid oxidation |
| 2 | Polyunsaturated fatty acid biosynthesis |
| 2 | De novo fatty acid biosynthesis |
| 2 | Phytanic acid peroxisomal oxidation |
| 2 | R Group Synthesis |
| 2 | Selenoamino acid metabolism |
| 3 | Phytanic acid peroxisomal oxidation |
| 3 | Omega-6 fatty acid metabolism |
| 4 | Glycosphingolipid biosynthesis - globoseries |
| 4 | Lysine metabolism |
| 4 | Tyrosine metabolism |
| 4 | Polyunsaturated fatty acid biosynthesis |
| 4 | Glycosphingolipid biosynthesis - ganglioseries |
| 4 | Keratan sulfate degradation |
| 4 | N-Glycan Degradation |
| 4 | Linoleate metabolism |
| 4 | Vitamin A (retinol) metabolism |
| 4 | Butanoate metabolism |
| 4 | Trihydroxycoprostanoyl-CoA beta-oxidation |
| 4 | Glycerophospholipid metabolism |
| 4 | Omega-3 fatty acid metabolism |
| 4 | Starch and Sucrose Metabolism |
| 5 | Purine metabolism |
| 5 | Leukotriene metabolism |
| 5 | Urea cycle/amino group metabolism |
| 5 | Methionine and cysteine metabolism |
| 5 | Tryptophan metabolism |
| 5 | Aminosugars metabolism |
| 5 | Aspartate and asparagine metabolism |

## 4. DISCUSSION

While altered metabolism is increasingly recognized as a crucial factor in the development of OA, further data are needed to understand the role of aberrant metabolism in OA pathophysiology. This study found distinct human cartilage-derived metabolomic profiles between healthy and end-stage OA patients. Through a comprehensive analysis aimed at discerning differences in the metabolome of healthy and OA cartilage, several metabolites and pathways associated with matrix metabolism, lipid metabolism, mitochondrial function, vitamin metabolism, and amino acid metabolism were differentially regulated between healthy and OA cartilage. Moreover, investigation of metabolic diversity within the metabolome of OA cartilage alone mapped to distinct metabolomic endotypes displaying the heterogeneous nature of OA. Considering these metabolomic findings, a greater understanding of altered cartilage metabolism in OA may lead to the identification of candidate biomarkers and drug targets to slow, halt, or reverse cartilage damage in end-stage OA.

### 4.1 Matrix metabolism

OA cartilage exhibited greater evidence of altered matrix metabolism compared to healthy cartilage. Specifically, keratan sulfate degradation and N-Glycan degradation were upregulated in OA cartilage compared to healthy cartilage. Keratan sulfate, a type of glycosaminoglycan (GAG), plays a vital role in cartilage matrix homeostasis and maintenance. GAG content in both synovial fluid (SF) and cartilage are indicative of joint health where an increase of GAGs in SF suggests increased cartilage turnover, which is subsequently reflected by a decrease in GAG content within the cartilage itself^25–27^. Furthermore, alterations in N-glycan degradation likely reflect changes in joint lubrication as N-glycans are an important component of lubricin, a glycoprotein that lines cartilage surface and serves as a key joint lubricant with chondroprotective properties^28^.

### 4.2 Lipid and mitochondria-related metabolism

Several lipid-related pathways were upregulated in OA cartilage compared to healthy cartilage and were differentially regulated across OA endotypes. Notably, the present study identified several significant lipid-related pathways that have been previously linked to OA, including the carnitine shuttle, arachidonic acid metabolism, omega-3 and −6 metabolism, glycosphingolipid metabolism, and glycerophospholipid metabolism. Cartilage relies on bone and SF for lipid transport, underscoring the critical role of lipid metabolism in maintaining cartilage homeostasis. Arachidonic acid (AA), leukotriene F4, N-acetyl-leukotriene E4, and panaxydol linoleate were identified using LC-MS/MS data and were at higher concentrations in OA cartilage compared to healthy and differed in abundance across OA endotypes (Supplemental Figures 1 and 3). AA, a type of omega-6 polyunsaturated fatty acid known to be associated with inflammation, is typically found in lower levels in healthy cartilage and increases as OA progresses^29^. Additionally, elevated AA have been detected in OA SF^16, 30^, synovium^31^, and more broadly, the severity of synovitis and histological changes in OA have been correlated with serum levels of omega-3 and −6^32, 33^.

The detection of perturbed lipid pathways and a handful of identified lipid species in OA cartilage may reflect adaptive responses in mitochondrial function and biofuel utilization in response to OA. While healthy cartilage relies on both glucose and lipids as energy sources, OA cartilage exhibits a greater dependence on lipids^34, 35^. This metabolic switch to lipid utilization can lead to the accumulation of lipids, increased production of reactive oxygen species and nitric oxide, decreased ATP production, leading to eventual tissue breakdown and death^36–39^.

Central to this metabolic switch is the carnitine shuttle, which plays a key role in regulating oxidative status by transporting lipids across the mitochondrial membrane to generate ATP. The upregulation of the carnitine shuttle in OA cartilage compared to healthy cartilage, and across OA endotypes, is supported by previous studies which not only detected the carnitine shuttle but also elevated levels of acylcarnitine and other carnitine species^17^ in SF of OA patients.

Moreover, cervonyl carnitine, a type of acylcarnitine, was identified using LC-MS/MS data and was significantly higher in all OA cartilage samples compared to healthy cartilage and differed in abundance across OA endotypes (Supplemental Figures 1 and 3). Cervonyl carnitine is often produced as a result of a disorder or disease (*i.e.*, cancer, diabetes, cardiovascular disease) and disrupts energy production^40^. It has been well documented that OA perturbs energy production in cartilage, therefore, the detection of this species in the present study could be a result of receiving cartilage from donors with radiography-confirmed OA.

To our knowledge, only one study to date has detected cervonyl carnitine in SF from patients who sustained a traumatic knee injury^18^. The authors hypothesized that a metabolic switch toward lipid utilization and the involvement of mechanisms like the carnitine shuttle were necessary to meet heightened energy demands post-injury. Furthermore, they speculated that ongoing analysis of these species may help manage post-traumatic OA. Thus, the detection of cervonyl carnitine in OA cartilage and in SF post-injury further highlights its potential as a marker that can be monitored over time to assess β-oxidation, joint heath, while also potentially predicting the onset and progression of OA. Furthermore, cervonyl carnitine warrants further investigation as a potential biomarker and druggable target for the purpose of slowing, halting, or reversing OA.

### 4.3 Vitamin metabolism

Vitamin E metabolism was notably upregulated in OA cartilage compared to healthy. Vitamin E has antioxidant properties, which could prove beneficial in counteracting the heightened oxidative stress experienced by the joint during OA^41^. Additionally, vitamin A was dysregulated across OA endotypes. The relationship between OA and vitamin A, including the Vitamin A derivative all-trans-retinoic acid, have garnered attention due to its key role in skeletal development and cartilage maintenance^42, 43^. Specifically, all-trans retinoic acid can regulate type X collagen and matrix metalloproteinase-13 driving a hypertrophic phenotype^43, 44^. Moreover, elevated vitamin A metabolite levels have been detected in SF, serum, and cartilage from OA individuals suggesting its potential role in OA within cartilage^42^.

In contrast, vitamin K metabolism was downregulated in OA cartilage compared to healthy cartilage. These findings align with prior research that explored the relationship between OA and vitamin K, which is important for its role as a cofactor for the carboxylation of vitamin K-dependent proteins, including matrix Gla proteins, osteocalcin, and Gas-6^45^. These proteins are present in the joint and play a key role in the maintenance of cartilage and bone. Their absence or deficiency can lead to an increased incidence and progression of knee OA^45–47^. Specifically, alterations in vitamin K levels parallel the abnormalities observed in OA disease progression, encompassing aspects such as hypertrophic and apoptotic chondrocytes, cartilage mineralization, and endochondral ossification^48, 49^.

### 4.4 Amino Acid Metabolism

Amino acid metabolism was significantly downregulated in OA cartilage compared to healthy cartilage. While histidine metabolism was not differently regulated across endotypes, the pronounced downregulation compared to healthy cartilage aligns with a previous study that identified declining trends in serum histidine levels as OA advances^50^. Additionally, the ratio of branched-chain amino acids to histidine has emerged as a potential indicator of disease progression^51^. In contrast, various amino acids including tryptophan, methionine, cysteine, aspartate, and asparagine were upregulated in healthy cartilage compared to OA cartilage confirmed by pairwise and endotype comparisons.

This pattern mirrors similar observations made in a study comparing SF metabolism from healthy, early, and late-stage OA patients, where these amino acid pathways were upregulated in healthy SF^16^. Focusing solely on OA cartilage, these same amino acid pathways displayed different regulation patterns across OA endotypes. This aligns with previous literature, indicating that these amino acids tend to decrease as OA progresses, with levels being highest in healthy cartilage, moderately high in early-stage OA, and diminishing with end-stage OA^52, 53^.

Furthermore, specific amino acids like glycine and alanine, both of which are abundant in collagen, have been putatively identified as potential markers to distinguish OA cartilage from healthy^7^. This observed dysregulation of amino acids may indicate their potential role in responding to disease and could reflect the degree of joint damage. Nevertheless, further research is required to underpin the relationship between amino acid metabolism and OA.

### 4.5 Limitations

While this study included healthy cartilage samples to examine disease-associated metabolic changes, it is not without limitations. Firstly, sample size for this study was not uniform where 11 healthy cartilage and 35 OA cartilage samples were obtained. Secondly, relevant clinical covariates (e.g., age, BMI, sex, prior medical history) were not available for the obtained healthy cartilage samples. Furthermore, age and sex were provided for all OA donors, but BMI was not provided. Considering partial information provided for both healthy and OA cartilage samples, age-, BMI-, and sex-matching analyses were not performed, nor can this information be used to shed light on driving factors that differentiate OA endotypes.

### 4.6 Conclusions

The results of this study provide clear evidence of OA-induced metabolic perturbations in human articular cartilage. Considering the heterogenous nature of OA, the detection of metabolic differences between healthy and OA individuals, and within OA individuals alone, can be further examined to pinpoint the diverse landscape of OA. With this approach, we uncovered specific metabolomic patterns and identified metabolites that may serve as valuable indicators of disease status or therapeutic targets. Expansion of this study will delineate joint-level metabolic activity in cartilage and how that is reflected, or associated, with the metabolism of other musculoskeletal tissues and fluids.

## ACKNOWLEDGEMENTS

Authors thank the Montana State University Mass Spectrometry Facility including Dr. Donald Smith and Jesse Thomas for assisting with mass spectrometry analysis, interpretation, and metabolite identification. Funding for the Mass Spectrometry Facility used in this publication was made possible by the M.J. Murdock Charitable Trust, the National Institute of General Medical Sciences of the National Institutes of Health (P20GM103474 and S10OD28650). Additionally, we thank Brady Hislop for his assistance in analyzing data and building data analysis pipelines. This study was funded by the National Institutes of Health under Award Numbers R01AR073964 and R01AR081489 (RKJ), the National Science Foundation under Award Number CMMI 1554708 (RKJ), the M.J. Murdock Charitable Trust under Award Numbers FSU-2017207 and NS-202016444 (AKH), and the National Aeronautics and Space Administration under Award Number 80NSSC20M0042 (AKH).

## Author Contributions

HDW, AKH, and RKJ designed experiments; HDW and PB harvested samples; HDW, AHW, ARB, EAH, MGG performed metabolite extractions; HDW ran LC-MS samples; HDW and AHW analyzed data; HDW, AHW, ARB, AKH, BB, and RKJ interpreted results; HDW, AHW, AKH, and RKJ drafted manuscript. All authors have read and revised the manuscript.

## Conflicts of Interest

Authors have no conflicts of interest to disclose. Dr. June owns stock in Beartooth Biotech and OpenBioWorks, which were not involved in this study.

---

**Supplemental Figure 1.**
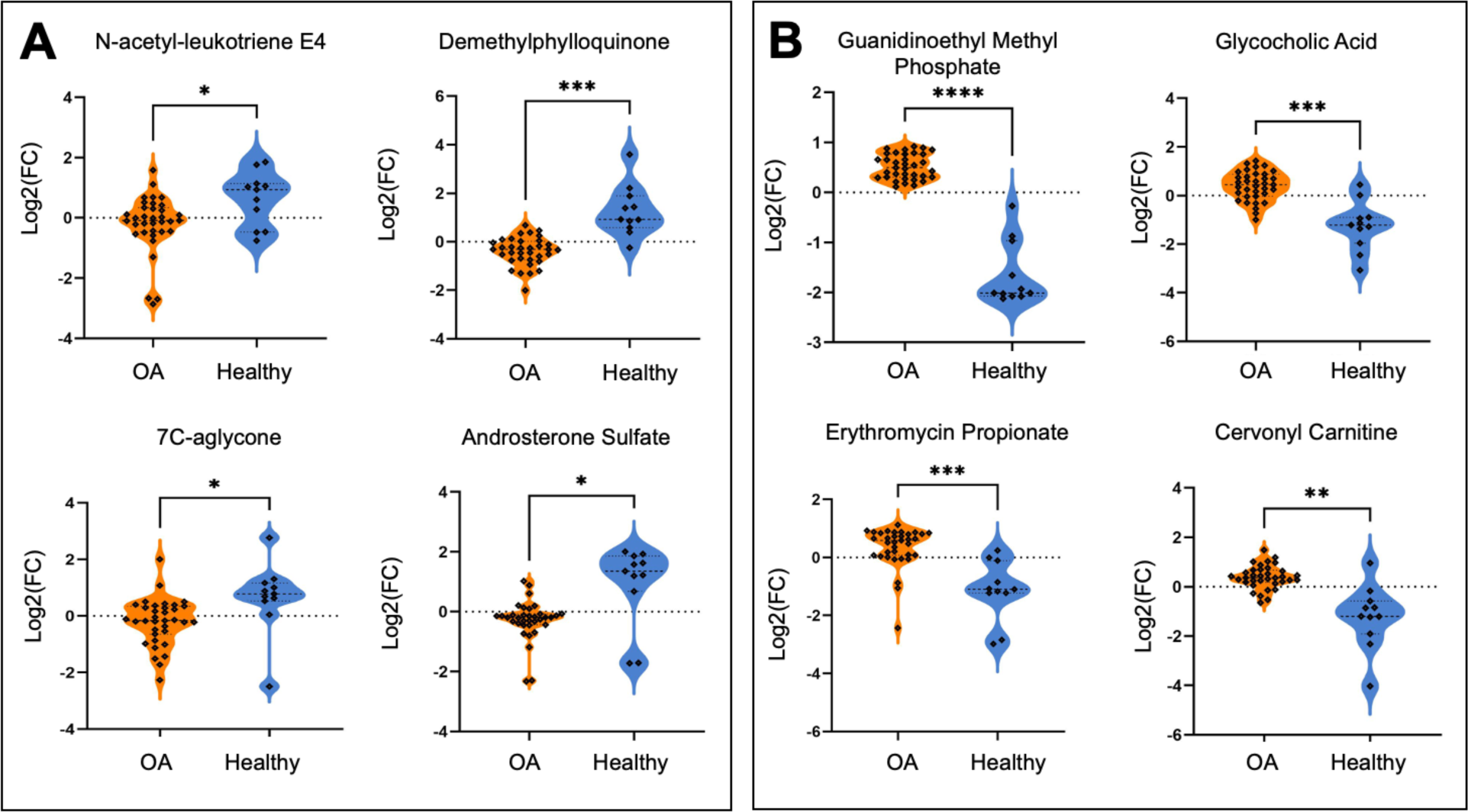
Identified metabolites differ in abundance between healthy and osteoarthritic cartilage. (A) Metabolites – N-acetyl-leukotriene E4, demethylphylloquinone, 7C-aglycone, androsterone sulfate – are higher in abundance in healthy cartilage compared to osteoarthritic cartilage. (B) Conversely, metabolites - guanidinoethyl methyl phosphate, glycocholic acid, erythromycin propionate, cervonyl carnitine – are higher in abundance in cartilage from individuals with osteoarthritis compared to healthy cartilage. Mass-to-charge intensities of interest were normalized and used to generate plots. To correct for multiple comparisons, FDR p-value corrections were performed and were less than < 0.05. Moreover, Welch’s t-tests were performed for each identified metabolite. Orange = osteoarthritis. Blue = healthy. **** = p < 0.0001, *** p < 0.0002, ** p < 0.001, * = p < 0.05.

**Supplemental Figure 2.**
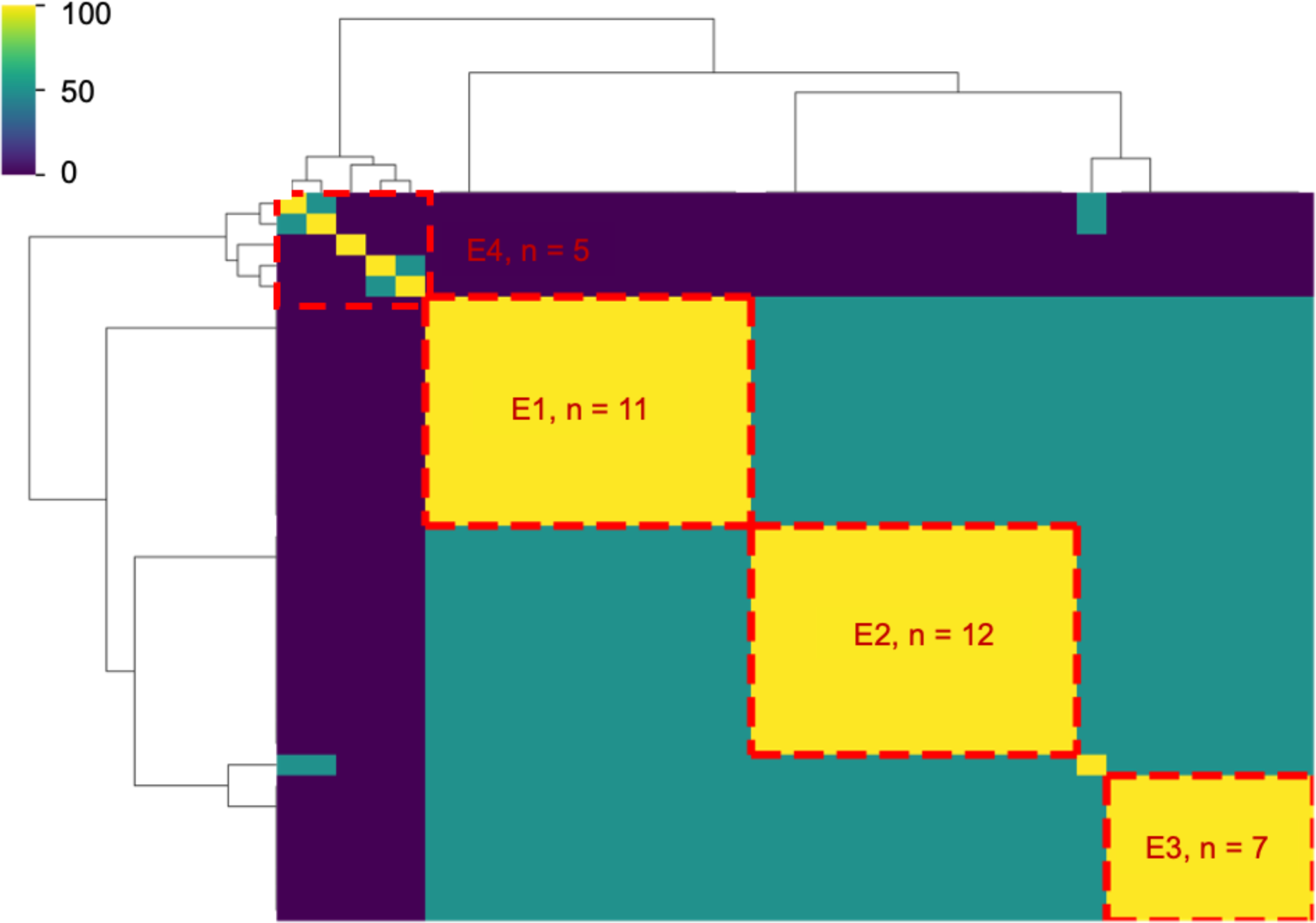
Ensemble clustering of osteoarthritis participants revealed distinct metabolic endotypes. By employing ensemble clustering of all osteoarthritis participants based on metabolomic data, four unique endotypes were revealed. Specifically, endotype one (E1) is composed of n = 11 participants. Moreover, endotype two (E2), three (E3), and four (E4) are composed of n = 12, n = 7, and n = 5, respectively. Scale bar represents the percentage (%) of time participants cluster together.

**Supplemental Figure 3.**
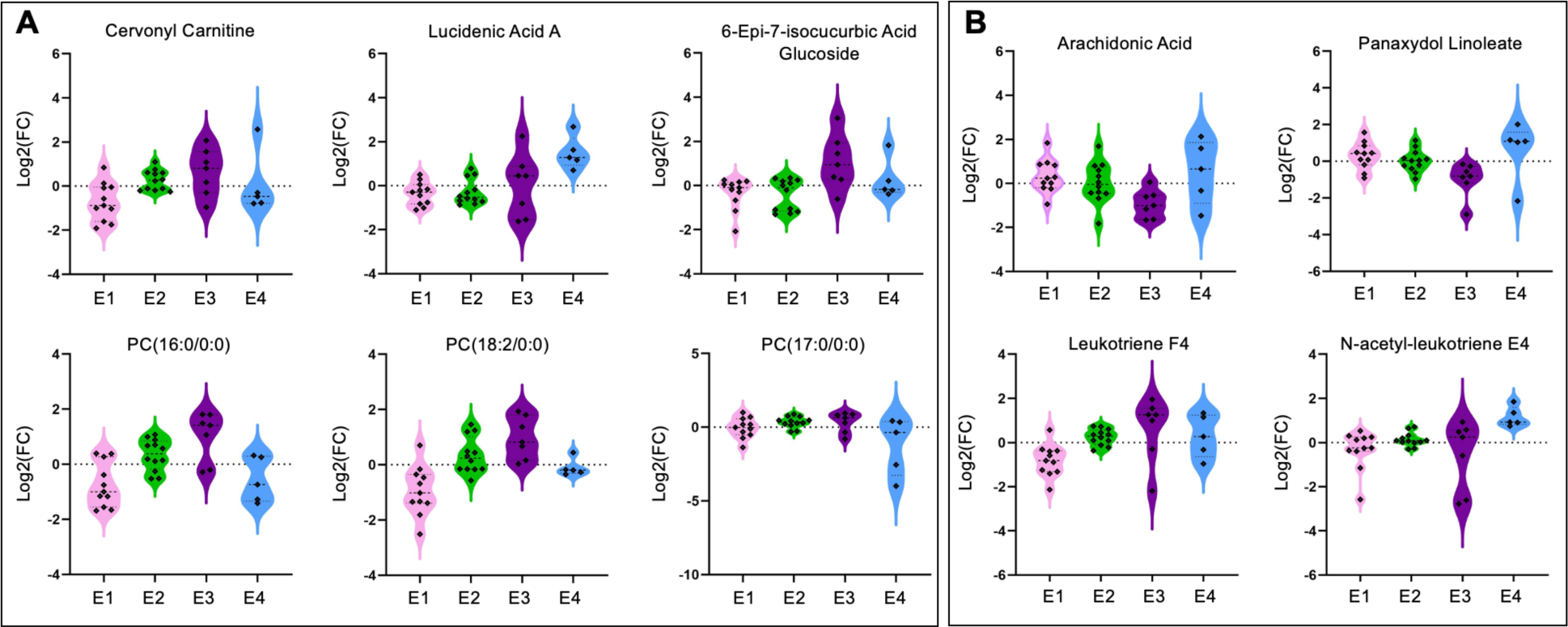
Identified metabolites differ in abundance across osteoarthritis endotypes. (A) Lipid and lipid-like identified metabolites that are differentially regulated across osteoarthritis endotypes. (B) Identified metabolites that differ in abundance across endotypes and are associated with arachidonic acid metabolism and leukotriene metabolism. Mass-to-charge intensities of interest were normalized and used to generate plots. To correct for multiple comparisons, FDR p-value corrections were performed and were less than < 0.05. Moreover, Welch’s t-tests were performed for each identified metabolite. Endotype colors correspond to: pink – Endotype 1; green – Endotype 2; purple – Endotype 3; blue – Endotype 4.

**Supplemental Table 1.** Participant information including donor number, assigned anonymous identifier, age, sex, and osteoarthritis endotype group.

**Supplemental Table 2.** All metabolic pathways determined from MetaboAnalyst when comparing healthy and diseased cartilage using volcano plot analyses. Populations of metabolite features are defined on Figure 1C.

**Supplemental Table 3.** Putatively identified metabolites that differ in abundance between healthy and diseased cartilage distinguished by volcano plot analysis. Identifications were made by performing liquid chromatography tandem mass spectrometry (LC-MS/MS). For all identified, information includes observed and theoretical mass-to-charge ratios, parts per million (ppm) error, accepted compound ID and description, adduct, chemical formula, total score out of 100, and fragmentation score. Identifications with error greater than 20 ppm, total score < 60, and a fragmentation score < 12 were excluded.

**Supplemental Table 4.** Putatively identified metabolites that differ in abundance between healthy and diseased cartilage distinguished by t-test analysis. Identifications were made by performing liquid chromatography tandem mass spectrometry (LC-MS/MS). For all identified, information includes observed and theoretical mass-to-charge ratios, parts per million (ppm) error, accepted compound ID and description, adduct, chemical formula, total score out of 100, and fragmentation score. Identifications with error greater than 20 ppm, total score < 60, and a fragmentation score < 12 were excluded.

**Supplemental Table 5.** Putatively identified metabolites that differ in abundance between osteoarthritis endotype groups distinguished by ANOVA analysis. Identifications were made by performing liquid chromatography tandem mass spectrometry (LC-MS/MS). For all identified, information includes observed and theoretical mass-to-charge ratios, parts per million (ppm) error, accepted compound ID and description, adduct, chemical formula, total score out of 100, and fragmentation score. Identifications with error greater than 20 ppm, total score < 60, and a fragmentation score < 12 were excluded. All metabolite features with an FDR-corrected p-value > 0.05 distinguished by ANOVA when comparing all four osteoarthritis groups.

**Supplemental Table 5.** All metabolic pathways determined from MetaboAnalyst when comparing osteoarthritis endotypes using median metabolite intensity heatmap analysis. Clusters defined on Figure 2E.

## Abbreviations List

OA: osteoarthritis
LC-MS: liquid chromatography-mass spectrometry
LC-MS/MS: liquid chromatography tandem mass spectrometry
HCA: hierarchical clustering analysis
PCA: principal component analysis
PLS-DA: partial least squares-discriminant analysis
FDR: false discovery rate
GAG: glycosaminoglycan
SF: synovial fluid
AA: arachidonic acid

## REFERENCES

1. Long H, Liu Q, Yin H, Wang K, Diao N, Zhang Y, et al. Prevalence trends of site-specific osteoarthritis from 1990 to 2019: findings from the Global Burden of Disease Study 2019. Arthritis Rheumatol 2022.

2. Vina ER, Kwoh CK. Epidemiology of osteoarthritis: literature update. Curr Opin Rheumatol 2018; 30: 160–167.

3. Barbour KE, Helmick CG, Boring M, Brady TJ. Vital Signs: Prevalence of Doctor-Diagnosed Arthritis and Arthritis-Attributable Activity Limitation - United States, 2013-2015. MMWR Morb Mortal Wkly Rep 2017; 66: 246-253.

4. Bitton R. The economic burden of osteoarthritis. Am J Manag Care 2009; 15: S230–235.

5. Hootman JM, Helmick CG, Barbour KE, Theis KA, Boring MA. Updated Projected Prevalence of Self-Reported Doctor-Diagnosed Arthritis and Arthritis-Attributable Activity Limitation Among US Adults, 2015-2040. Arthritis Rheumatol 2016; 68: 1582-1587.

6. Neogi T. The epidemiology and impact of pain in osteoarthritis. Osteoarthritis Cartilage 2013; 21: 1145–1153.

7. Shet K, Siddiqui SM, Yoshihara H, Kurhanewicz J, Ries M, Li X. High-resolution magic angle spinning NMR spectroscopy of human osteoarthritic cartilage. NMR Biomed 2012; 25: 538–544.

8. Xue M, Huang N, Luo Y, Yang X, Wang Y, Fang M. Combined Transcriptomics and Metabolomics Identify Regulatory Mechanisms of Porcine Vertebral Chondrocyte Development In Vitro. Int J Mol Sci 2024; 25.

9. Zignego DL, Hilmer JK, June RK. Mechanotransduction in primary human osteoarthritic chondrocytes is mediated by metabolism of energy, lipids, and amino acids. J Biomech 2015; 48: 4253–4261.

10. Bartlett SJ, Ling SM, Mayo NE, Scott SC, Bingham CO, 3rd. Identifying common trajectories of joint space narrowing over two years in knee osteoarthritis. Arthritis Care Res (Hoboken) 2011; 63: 1722–1728.

11. Collins JE, Katz JN, Dervan EE, Losina E. Trajectories and risk profiles of pain in persons with radiographic, symptomatic knee osteoarthritis: data from the osteoarthritis initiative. Osteoarthritis Cartilage 2014; 22: 622–630.

12. Karsdal MA, Bihlet A, Byrjalsen I, Alexandersen P, Ladel C, Michaels M, et al. OA phenotypes, rather than disease stage, drive structural progression--identification of structural progressors from 2 phase III randomized clinical studies with symptomatic knee OA. Osteoarthritis Cartilage 2015; 23: 550–558.

13. Bruyere O, Cooper C, Arden N, Branco J, Brandi ML, Herrero-Beaumont G, et al. Can we identify patients with high risk of osteoarthritis progression who will respond to treatment? A focus on epidemiology and phenotype of osteoarthritis. Drugs Aging 2015; 32: 179–187.

14. Deveza LA, Nelson AE, Loeser RF. Phenotypes of osteoarthritis: current state and future implications. Clin Exp Rheumatol 2019; 37 Suppl 120: 64–72.

15. Patti GJ, Yanes O, Siuzdak G. Innovation: Metabolomics: the apogee of the omics trilogy. Nat Rev Mol Cell Biol 2012; 13: 263–269.

16. Carlson AK, Rawle RA, Wallace CW, Brooks EG, Adams E, Greenwood MC, et al. Characterization of synovial fluid metabolomic phenotypes of cartilage morphological changes associated with osteoarthritis. Osteoarthritis Cartilage 2019; 27: 1174–1184.

17. Zhang W, Likhodii S, Zhang Y, Aref-Eshghi E, Harper PE, Randell E, et al. Classification of osteoarthritis phenotypes by metabolomics analysis. BMJ Open 2014; 4: e006286.

18. Welhaven HD, Welfley AH, Pershad P, Satalich J, O’Connell R, Bothner B, et al. Metabolic phenotypes reflect patient sex and injury status: A cross-sectional analysis of human synovial fluid. Osteoarthritis Cartilage 2023.

19. Kessner D, Chambers M, Burke R, Agus D, Mallick P. ProteoWizard: open source software for rapid proteomics tools development. Bioinformatics 2008; 24: 2534–2536.

20. Smith CA, Want EJ, O’Maille G, Abagyan R, Siuzdak G. XCMS: processing mass spectrometry data for metabolite profiling using nonlinear peak alignment, matching, and identification. Anal Chem 2006; 78: 779–787.

21. Welhaven HD, Vahidi G et al. The Cortical Bone Metabolome of C57BL/6J Mice Is Sexually Dimorphic. JBMR Plus 2022; 6.

22. Pang Z, Zhou G, Ewald J, Chang L, Hacariz O, Basu N, et al. Using MetaboAnalyst 5.0 for LC-HRMS spectra processing, multi-omics integration and covariate adjustment of global metabolomics data. Nat Protoc 2022; 17: 1735–1761.

23. Xiao JF, Zhou B, Ressom HW. Metabolite identification and quantitation in LC-MS/MS-based metabolomics. Trends Analyt Chem 2012; 32: 1–14.

24. Wishart DS, Guo A, Oler E, Wang F, Anjum A, Peters H, et al. HMDB 5.0: the Human Metabolome Database for 2022. Nucleic Acids Res 2022; 50: D622–D631.

25. Elliott RJ, Gardner DL. Changes with age in the glycosaminoglycans of human articular cartilage. Ann Rheum Dis 1979; 38: 371–377.

26. Hjertquist SO, Lemperg R. Identification and concentration of the glycosaminoglycans of human articular cartilage in relation to age and osteoarthritis. Calcif Tissue Res 1972; 10: 223–237.

27. Thonar EJ, Masuda K, Hauselmann HJ, Uebelhart D, Lenz ME, Manicourt DH. Keratan sulfate in body fluids in joint disease. Acta Orthop Scand Suppl 1995; 266: 103–106.

28. Jay GD, Torres JR, Warman ML, Laderer MC, Breuer KS. The role of lubricin in the mechanical behavior of synovial fluid. Proc Natl Acad Sci U S A 2007; 104: 6194–6199.

29. Ioan-Facsinay A, Kloppenburg M. Bioactive lipids in osteoarthritis: risk or benefit? Curr Opin Rheumatol 2018; 30: 108–113.

30. Van de Vyver A, Clockaerts S, van de Lest CHA, Wei W, Verhaar J, Van Osch G, et al. Synovial Fluid Fatty Acid Profiles Differ between Osteoarthritis and Healthy Patients. Cartilage 2020; 11: 473–478.

31. Cillero-Pastor B, Eijkel G, Kiss A, Blanco FJ, Heeren RM. Time-of-flight secondary ion mass spectrometry-based molecular distribution distinguishing healthy and osteoarthritic human cartilage. Anal Chem 2012; 84: 8909–8916.

32. Baker KR, Matthan NR, Lichtenstein AH, Niu J, Guermazi A, Roemer F, et al. Association of plasma n-6 and n-3 polyunsaturated fatty acids with synovitis in the knee: the MOST study. Osteoarthritis Cartilage 2012; 20: 382–387.

33. Lippiello L, Walsh T, Fienhold M. The association of lipid abnormalities with tissue pathology in human osteoarthritic articular cartilage. Metabolism 1991; 40: 571–576.

34. Dalmao-Fernandez A, Lund J, Hermida-Gomez T, Vazquez-Mosquera ME, Rego-Perez I, Blanco FJ, et al. Impaired Metabolic Flexibility in the Osteoarthritis Process: A Study on Transmitochondrial Cybrids. Cells 2020; 9.

35. Smith RL, Soeters MR, Wust RCI, Houtkooper RH. Metabolic Flexibility as an Adaptation to Energy Resources and Requirements in Health and Disease. Endocr Rev 2018; 39: 489–517.

36. Blanco FJ, Lopez-Armada MJ, Maneiro E. Mitochondrial dysfunction in osteoarthritis. Mitochondrion 2004; 4: 715–728.

37. Blanco FJ, Rego I, Ruiz-Romero C. The role of mitochondria in osteoarthritis. Nat Rev Rheumatol 2011; 7: 161–169.

38. Lane RS, Fu Y, Matsuzaki S, Kinter M, Humphries KM, Griffin TM. Mitochondrial respiration and redox coupling in articular chondrocytes. Arthritis Res Ther 2015; 17: 54.

39. Wu L, Liu H, Li L, Liu H, Cheng Q, Li H, et al. Mitochondrial pathology in osteoarthritic chondrocytes. Curr Drug Targets 2014; 15: 710–719.

40. Dambrova M, Makrecka-Kuka M, Kuka J, Vilskersts R, Nordberg D, Attwood MM, et al. Acylcarnitines: Nomenclature, Biomarkers, Therapeutic Potential, Drug Targets, and Clinical Trials. Pharmacol Rev 2022; 74: 506–551.

41. Collins JA, Wood ST, Nelson KJ, Rowe MA, Carlson CS, Chubinskaya S, et al. Oxidative Stress Promotes Peroxiredoxin Hyperoxidation and Attenuates Pro-survival Signaling in Aging Chondrocytes. J Biol Chem 2016; 291: 6641–6654.

42. Davies MR, Ribeiro LR, Downey-Jones M, Needham MR, Oakley C, Wardale J. Ligands for retinoic acid receptors are elevated in osteoarthritis and may contribute to pathologic processes in the osteoarthritic joint. Arthritis Rheum 2009; 60: 1722–1732.

43. Underhill TM, Weston AD. Retinoids and their receptors in skeletal development. Microsc Res Tech 1998; 43: 137–155.

44. Flannery CR, Little CB, Caterson B, Hughes CE. Effects of culture conditions and exposure to catabolic stimulators (IL-1 and retinoic acid) on the expression of matrix metalloproteinases (MMPs) and disintegrin metalloproteinases (ADAMs) by articular cartilage chondrocytes. Matrix Biol 1999; 18: 225–237.

45. Misra D, Booth SL, Tolstykh I, Felson DT, Nevitt MC, Lewis CE, et al. Vitamin K deficiency is associated with incident knee osteoarthritis. Am J Med 2013; 126: 243–248.

46. Shea MK, Kritchevsky SB, Hsu FC, Nevitt M, Booth SL, Kwoh CK, et al. The association between vitamin K status and knee osteoarthritis features in older adults: the Health, Aging and Body Composition Study. Osteoarthritis Cartilage 2015; 23: 370–378.

47. Wallin R, Schurgers LJ, Loeser RF. Biosynthesis of the vitamin K-dependent matrix Gla protein (MGP) in chondrocytes: a fetuin-MGP protein complex is assembled in vesicles shed from normal but not from osteoarthritic chondrocytes. Osteoarthritis Cartilage 2010; 18: 1096–1103.

48. Luo G, Ducy P, McKee MD, Pinero GJ, Loyer E, Behringer RR, et al. Spontaneous calcification of arteries and cartilage in mice lacking matrix GLA protein. Nature 1997; 386: 78–81.

49. Price PA, Williamson MK, Haba T, Dell RB, Jee WS. Excessive mineralization with growth plate closure in rats on chronic warfarin treatment. Proc Natl Acad Sci U S A 1982; 79: 7734–7738.

50. Zhang Q, Li H, Zhang Z, Yang F, Chen J. Serum metabolites as potential biomarkers for diagnosis of knee osteoarthritis. Dis Markers 2015; 2015: 684794.

51. Zhai G, Wang-Sattler R, Hart DJ, Arden NK, Hakim AJ, Illig T, et al. Serum branched-chain amino acid to histidine ratio: a novel metabolomic biomarker of knee osteoarthritis. Ann Rheum Dis 2010; 69: 1227–1231.

52. Abdelrazig S, Ortori CA, Doherty M, Valdes AM, Chapman V, Barrett DA. Metabolic signatures of osteoarthritis in urine using liquid chromatography-high resolution tandem mass spectrometry. Metabolomics 2021; 17: 29.

53. Igari T, Tsuchizawa M, Shimamura T. Alteration of tryptophan metabolism in the synovial fluid of patients with rheumatoid arthritis and osteoarthritis. Tohoku J Exp Med 1987; 153: 79–86.

