## Supplemental Figures for "Metabolomic Profiles and Pathways in Osteoarthritic Human Cartilage: A Comparative Analysis with Healthy Cartilage"

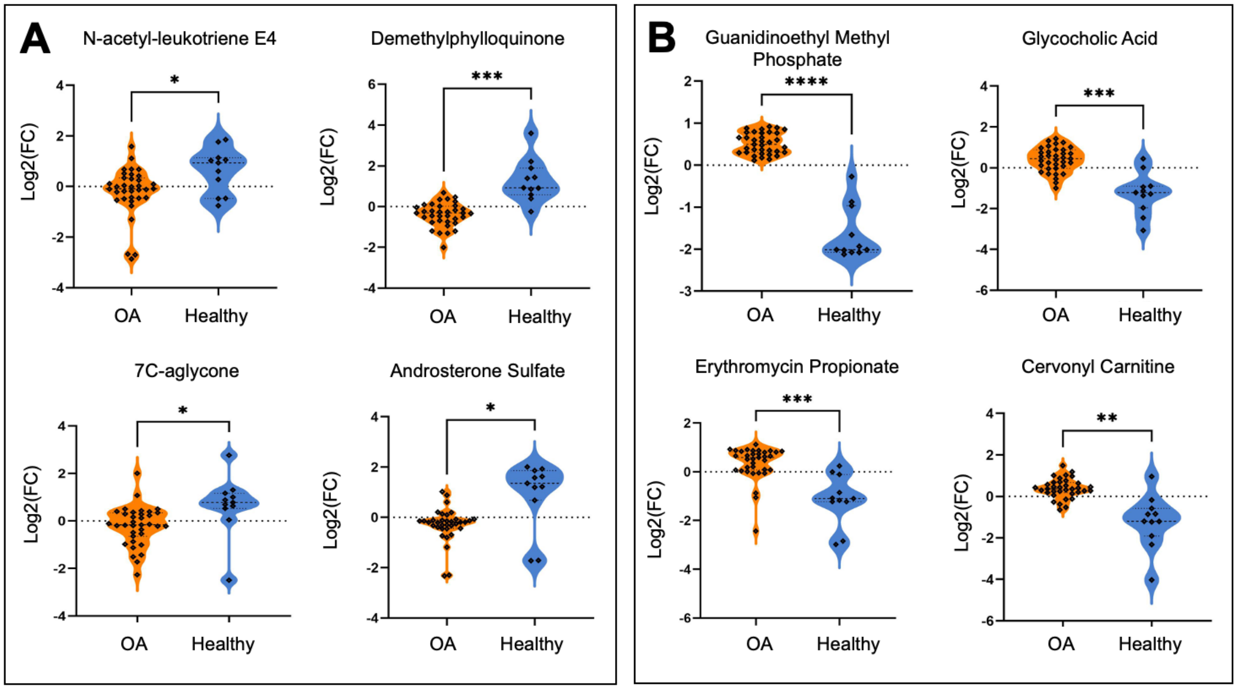


**Supplemental Figure 1. Identified metabolites differ in abundance between healthy and osteoarthritic cartilage.** (A) Metabolites – N-acetyl-leukotriene E4, demethylphylloquinone, 7C-aglycone, androsterone sulfate – are higher in abundance in healthy cartilage compared to osteoarthritic cartilage. (B) Conversely, metabolites - guanidinoethyl methyl phosphate, glycocholic acid, erythromycin propionate, cervonyl carnitine – are higher in abundance in cartilage from individuals with osteoarthritis compared to healthy cartilage. Mass-to-charge intensities of interest were normalized and used to generate plots. To correct for multiple comparisons, FDR p-value corrections were performed and were less than < 0.05. Moreover, Welch's t-tests were performed for each identified metabolite. Orange = osteoarthritis. Blue = healthy. **** = p < 0.0001, *** p < 0.0002, ** p < 0.001, * = p < 0.05.

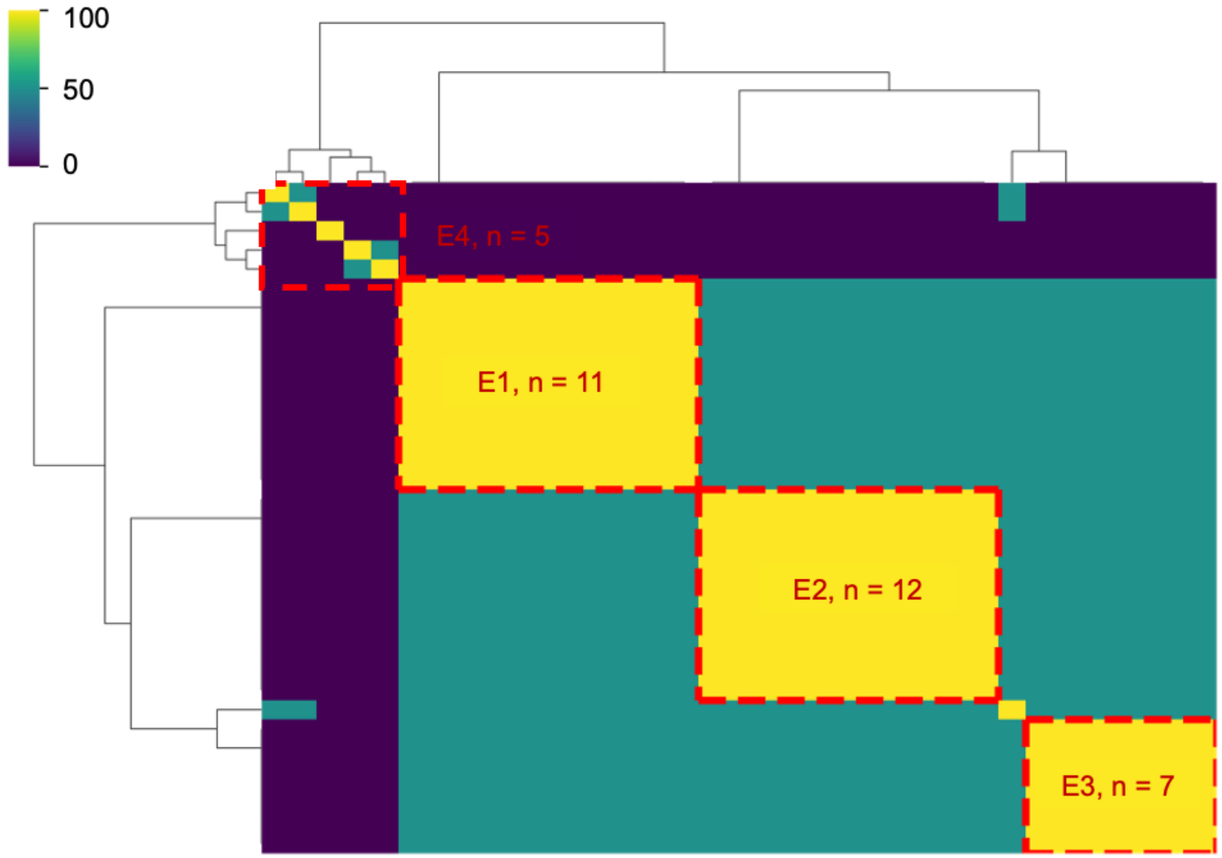


**Supplemental Figure 2. Ensemble clustering of osteoarthritis participants revealed distinct metabolic endotypes.** By employing ensemble clustering of all osteoarthritis participants based on metabolomic data, four unique endotypes were revealed. Specifically, endotype one (E1) is composed of n = 11 participants. Moreover, endotype two (E2), three (E3), and four (E4) are composed of n = 12, n = 7, and n = 5, respectively. Scale bar represents the percentage (%) of time participants cluster together.

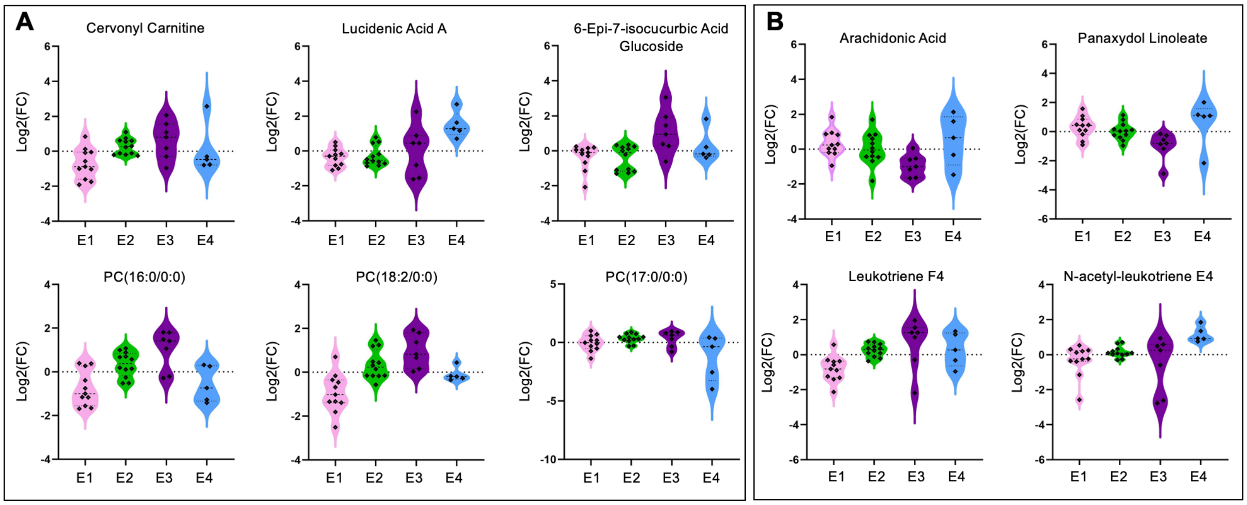


**Supplemental Figure 3. Identified metabolites differ in abundance across osteoarthritis endotypes.** (A) Lipid and lipid-like identified metabolites that are differentially regulated across osteoarthritis endotypes. (B) Identified metabolites that differ in abundance across endotypes and are associated with arachidonic acid metabolism and leukotriene metabolism. Mass-to-charge intensities of interest were normalized and used to generate plots. To correct for multiple comparisons, FDR p-value corrections were performed and were less than < 0.05. Moreover, Welch's t-tests were performed for each identified metabolite. Endotype colors correspond to: pink – Endotype 1; green – Endotype 2; purple – Endotype 3; blue – Endotype 4.
